# LabNet: hardware control software for the Raspberry Pi

**DOI:** 10.1101/2022.02.25.481971

**Authors:** Alexej Schatz, York Winter

**Author notes:** **For correspondence:** (AS); (YW).

## Abstract

Single-board computers such as the Raspberry Pi make it easy to control hardware setups for laboratory experiments. GPIOs and expansion boards (HATs) give access to a whole range of sensor and control hardware. However, controlling such hardware can be challenging, when many experimental setups run in parallel and the time component is critical. LabNet is a C++ optimized control layer software to give access to the Raspberry Pi connected hardware over a simple network protocol. LabNet was developed to be suitable for time-critical operations, and to be simple to expand. It leverages the actor model to simplify multithreading programming and to increase modularity. The message protocol is implemented in Protobuf and offers performance, small message size, and supports a large number of programming languages on the client side. It shows good performance compared to locally executed tools like Bpod or pyControl, outperforms Autopilot and reaches sub-millisecond range in network communication latencies. LabNet itself does not provide support for the design of experimental tasks. This is left to the client. LabNet can be used for general automation in experimental laboratories with its control PC located at some distance. LabNet is open source and under continuing development.

## Introduction

The combination of open source software, low cost microcontroller electronics, and the easy access to digital fabrication have led to a plethora of open source solutions for animal behaviour experimental systems (Open Behaviour (***Laubach et al., 2021***), Bpod (***Sanders, 2021***), Autopilot (***Saunders and Wehr, 2019***), pyControl (***Akam et al., 2022***), MiniScope (***Cai et al., 2016***), Bonsai (***Lopes et al., 2015***), Whisker (***Cardinal and Aitken, 2010***), OpenEphys GUI (***Siegle et al., 2017***)). Using our 10-year experience with the Rasperry Pi for animal behaviour experimental control and after two decades with different self-developed embedded control approaches we have developed in C++ a new, powerful, and highly versatile platform for hardware control via the Raspberry Pi.

We had two major goals: the platform has to be suitable for time-critical operations and be easy to extend. Furthermore, LabNet had to support a wide variety of hardware components and simultaneously control multiple animal behaviour experimental operant boxes. When conducting automated behavioural experiments, it is advantageous to test many animals in parallel with identical or, if necessary, individually-specific experiments. This is the only way to obtain complete data sets quickly and can only be achieved through automation. Figure 1 shows examples of operant conditioning cages (Skinner boxes (***Skinner, 1938***)) as controlled by LabNet. Our intention was not to create a completely new ecosystem like Autopilot. We wanted to simplify communication with hardware for projects using their coding language of choice on the PC. Also, we wanted to remain general so that LabNet can become a general platform for experimental laboratory automation.

**Figure 1.**
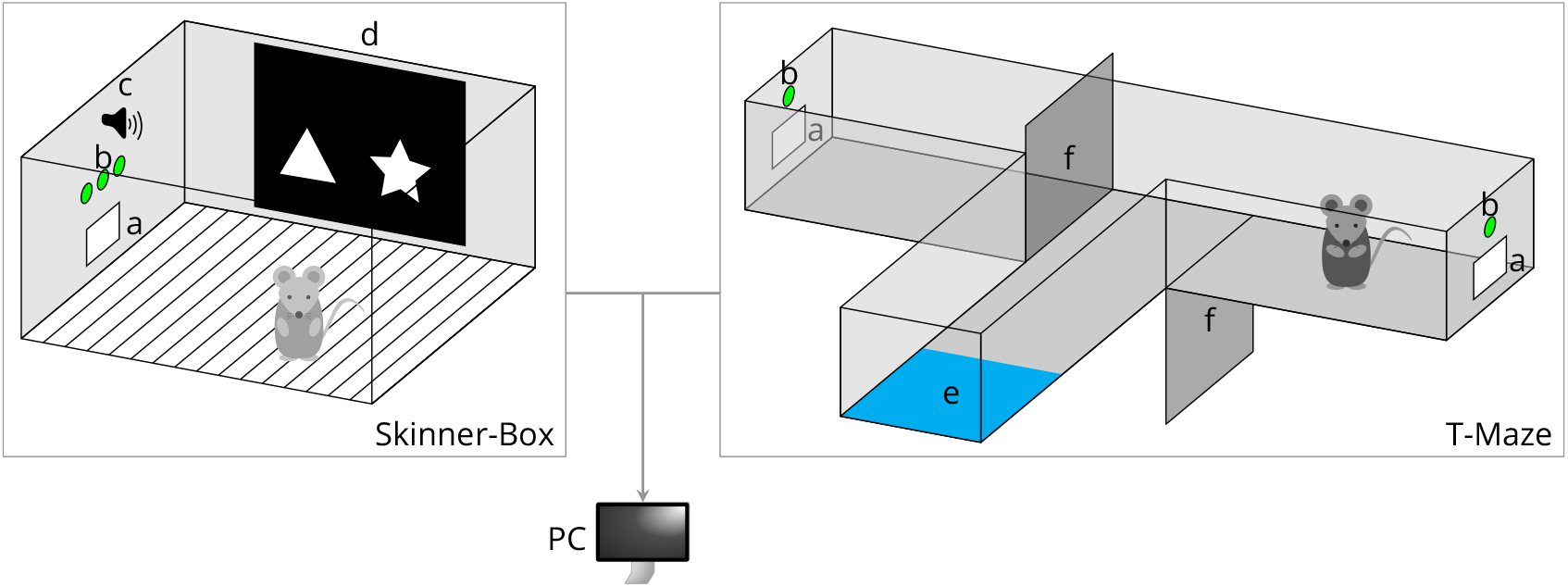
Examples of behavoiural set-ups controlled by LabNet. A Skinner box (left) contains (a) a feeder magazine that typically has a photo gate for nose-poke detection and a reward pellet dispenser. It also has (b) a row of LEDs and (c) a tone generator. (d) A monitor displays visual stimuli and may have a touch sensor for touchscreen functionality. The T-Maze (right) also has (a) a food magazine and (b) LEDs, and furthermore (e) an optical sensor to detect the return of the mouse to the start position and (f) two motorized doors that can be lowered to restrict access to the arms. ^1^

We selected the Raspberry Pi because it is low cost, powerful, has a wide selection of I/O addons and software components. To keep signal lines short, we gave each experimental setup its own Raspberry Pi. All systems are connected to the Ethernet network and are manageable via a central instance. This instance can be a normal PC and be located outside of the laboratory. Thus, the condition of experimental states, and in our case the animals, can be monitored at any time, even without entering the laboratory.

Autopilot also uses a swarm of Raspberry Pis. However, it implements a hierarchy where each Raspberry Pi can take a different role. This requires an additional configuration step and can complicate troubleshooting. We wanted to avoid this as well. This is the reason why each Raspberry Pi should run the same software and overall experimental control is executed by the central instance that is for example run from a PC. This separation also determined the network architecture of the entire system. The local instances on the Raspberry Pi are servers, and the central instance is the client. There can be one client to control experiments on all systems or multiple clients each controlling an experiment on one or more systems.

In the Performance Evaluation section, we compare the performance of LabNet to other solutions such as Autopilot, Whisker, Bpod and pyControl. The Design section outlines the general design of LabNet and explains the reasoning behind our choice of architecture. The Implementation section then provides some implementation details to compare LabNet with other systems in Related work.

## Design

Since the Raspberry Pi is a single-board computer, it runs “Raspberry Pi OS”: a Debian based Linux distribution. This allows a large freedom in the choice of programming language and software tools. Both interpreted languages, such as Python, and compiled languages are available. LabNet was required to meet two criteria:

1. time-critical: all operations should be performed as quickly as possible;
2. flexible: new functionality extensions should be as simple as possible.

Unfortunately, all interpreted programming languages have a disadvantage in execution speed compared to compiled languages. Nevertheless, many of the tools developed recently, such as Autopilot and pyControl, use Python. Python is a simple language and provides many packages for all purposes. However, because execution speed was of primary importance we decided to use C++.

Since its version 2 the Raspberry Pi has 4 cores additionally most hardware controllers, such as USB or ethernet operate asynchronously. LabNet needed an architecture that optimally leverages this already available hardware asynchrony for parallel execution. Handling GPIO lines is rapid, but accessing a UART may lead to considerable delays in sequential execution. This presupposes the use of multiple threads. Since programming with many parallel threads is an error-prone and time-consuming task, we decided to develop an actor-based software. This provides also higher flexibility and software modularity. Furthermore, a flexible and fast message protocol was also required for communication with the outside world (section Message protocol).

### Actor model

Developing a system with multiple threads still requires much care and can be challenging. Thread local state and program global state have to be protected. Some type of locking mechanism is required. Unfortunately, the locking mechanism itself increases not only the scalability but also the code complexity and error-proneness due to the locking order. Locking problems such as race conditions or dead-locks must be avoided. But time and execution order dependent errors can be difficult to find and fix.

LabNet is a concurrent system. The operations on the GPIOs, sending and receiving data via UARTs, sound output, etc. has to be independent from each other. For such purpose, message-passing approaches have been developed. In those, all inter-thread state sharing is encapsulated within messages sent between threads. All messages must be immutable or be copied for each thread.

Hewitt, Bishop and Steiger (***Hewitt et al., 1973***) proposed in 1973 one of the first message-passing systems in their actor model. Actors are active objects that communicate only over messages. Each actor has only knowledge about itself and his own functioning (shared-nothing principle). No global state exists in an actor system. Messages also do not block the sender (fire-and-forget principle). Hoare (***Hoare, 1978***) later introduced communicating sequential processes (CSP). The processes in CSP run asynchronously. They send messages that request or provide a piece of information. (Pre)Defined primitives encourage certain communication constructs and patterns, e.g. interleave results among many processes or the waiting for one of many processes to send data of interest.

These ideas have raised the level of abstraction from only considering shared memory to independent actors that communicate through a well-defined message protocol. In the late 80s, Ericsson developed Erlang (***Armstrong, 1996***), an actor-based programming language, and successfully used it in ATM network switches. The programming language Go from Google (***Google, 2021a***) leverages the CSP idea for concurrency. The Akka (***Lightbend, 2021***) framework was released in 2009 for Java and Scala.

### SObjectizer

From the several actor model libraries that are available for C++ such as the C++ Actor Framework (CAF) (***Charousset et al., 2013***), SObjectizer (***Stiffstream, 2021***), or Theron (***Mason, 2019***) we chose SObjectizer.

In SObjectizer a class or struct is sufficient to define a message. Actors are also normal classes derived from an `agent_t` base class. Thus, actors automatically have a “message box” (Mbox), through which messages can be received, and also methods that are automatically called, e.g. before an actor is started or stopped.

An Mbox can receive messages of all possible types. The Mbox of an actor has no name and must be communicated to other actors. However, named Mboxes can also exist. A reference to such an Mbox can be created anytime via its name. This is practical for accessing actors that always exist.

It is also possible to mix actors with other paradigms. This allows to move some parts of the application into the Boost ASIO or into threads. Mboxes can still be used for communication with actors from the outside. For communication with code from outside the actor world, so-called MChains are used. An MChain looks like a queue: actors can place messages there and threads can pull them at a later time.

One important feature of SObjectizer is the built-in support for hierarchical state machines (HSMs). All actors in SObjectizer are state machines. They can pass through several states in their lives and react to incoming messages depending on their current state.

Dispatchers are another important cornerstone of SObjectizer. Dispatchers provide an actor with the working context. They manage all message queues and execute the actors. Dispatcher types are:

1. `one_thread`: all actors are executed inside one thread. No true parallel execution;
2. `active_obj`: each actor is executed in its own thread. Maximum parallel execution;
3. `thread_pool`: this dispatcher creates a pool of worker threads. It provides a good compromise between thread management overhead and parallelism.

For our purposes the `thread_pool` is sufficient. But it is still possible to use other dispatcher types without having to adapt the actors.

### Message protocol

Our criteria for choosing the serialization tool were good performance, small message size and support in as many programming languages as possible. Text-based serialization formats, such as XML, JSON or ASCII-based plain-text have the advantage of being human-readable. The Whisker (***Cardinal and Aitken, 2010***) server uses an ASCII-based format. The disadvantages are the message size, higher computing requirements and, at least for the ASCII version, a custom message parser.

For our application, a binary format is a better solution, and we chose Protobuf (***Google, 2021b***) from Google. It is very popular and offers support for many programming languages. However, Protobuf has some disadvantages. For example, it is not the most memory or computationally efficient tool. Libraries such as Flatbuffers, Cap’n Proto or Simple Binary Encoding (SBE) are more efficient. However, these negative aspects of Protobuf only become critical when sending extremely large messages or at a very high rate. This is typically not the case in experiments that focus on actions of animal behaviour.

Protobuf uses a special meta-language to define messages. With `protoc`-generator, it is possible to create these message protocols for each supported programming language. Files with message definitions are a part of the Git repository.

One Protobuf disadvantage must be mentioned. A serialized Protobuf message contains no information about the byte length nor the message type. Protobuf leaves this information to the transmission medium. We have solved this simply: each message has at the beginning the message type and size. Both pieces of information are encoded as a number in Protobuf’s `varint` notation and are easy to parse with the Protobuf API.

## Implementation

The current implementation does not contain configuration files for LabNet. The hardware initialization is exclusively performed through client messages. LabNet comprises several loosely coupled actors. The most important are briefly described below (see also Figure 2).

**Figure 2.**
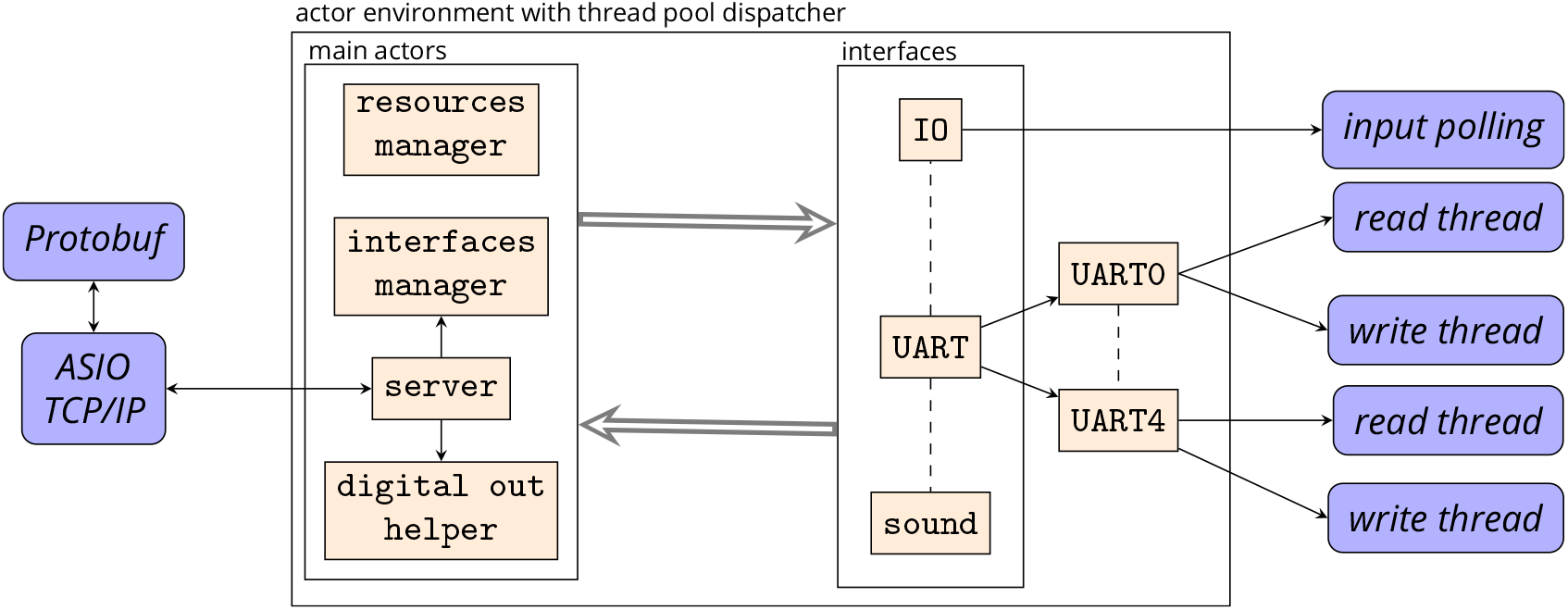
The core of LabNet is the actor environment with thread pool dispatcher. Within the environment, the main actors are always present. The actors of the individual interfaces are started by the interfaces manager as needed. The interface actors can also outsource their work to other actors and threads. All main and interface actors can communicate with each other. The threads themselves and network communication via Boost ASIO, on the other hand, are hidden behind their actors.

The network communication runs over TCP/IP. LabNet is the server. The current implementation accepts only one client connection. The main reason for this: multiple clients would require an additional resource management in LabNet so that no client can access foreign hardware. The server is implemented in Boost ASIO (***Boost, 2021***). The implementation is hidden behind the `server` actor. This actor can send and receive the Protobuf messages and also informs the actor world about the connection state. If the connection is lost, the actors can stop their work and resume it automatically later on a reconnection.

At the beginning, no single interface actor to communicate with the hardware exists. They are automatically created and started by the `interface manager`. Currently, several “interfaces” exist for controlling the GPIOs in different pin numbering schemes. Although WiringPi (***Henderson, 2019***) is used internally, BCM numbers can also be used. Furthermore, data can be received and sent via the internal UART and UART converters connected to the USB ports. Moreover, a simple sound output in the form of sine tones over HDMI or a headphone jack is possible.

Many pins on the Raspberry Pi offer more than one functionality. Clear responsibility for a hardware resource must be ensured. Each “interface” actor must request the resources from the `resource manager`. This is one of the first steps during the actor initialization phase.

The “interfaces” with digital outputs offer only the possibility to switch the output pin state. More complex procedures are implemented by the `digital out helper` actor. This actor can automatically turn off a pin after a defined time or generate pulses by specifying the on/off duration and number of pulses. Additionally, a group of pins can be automatically switched on and off together in a loop.

### Example

The following example shows the use of some of the LabNet messages. In this example, communication via TCP/IP is omitted for simplicity. We simply assume that a TCP/IP client exists and handles all operations like send, receive and serialization.

Let us assume an experimental setup with an LED and audio as stimuli, a valve to release a liquid reward and a photo gate as a nose-poke sensor to detect animal behaviour. All these components can be connected directly to the GPIO pins via a simple circuit. The headphone jack can be used for audio output. Then we need to send five commands to LabNet to initialize everything; see Listing 1. It would usually be necessary to wait for the response from LabNet and check the initialization result. Here, we skip this step.

During experiments, animals must usually perform some operant behaviour. This can be as simple as nose poking to trigger a photo gate after a certain stimulus has been perceived. In Listing 2, LabNet switches an LED and produces a sine tone. In the case of the tone, it is instructed to automatically generate a pulsed output. LabNet transmits on detection a photo gate’s state change to the client, both on and off. In response to a photo gate state change, a reward can be provided.

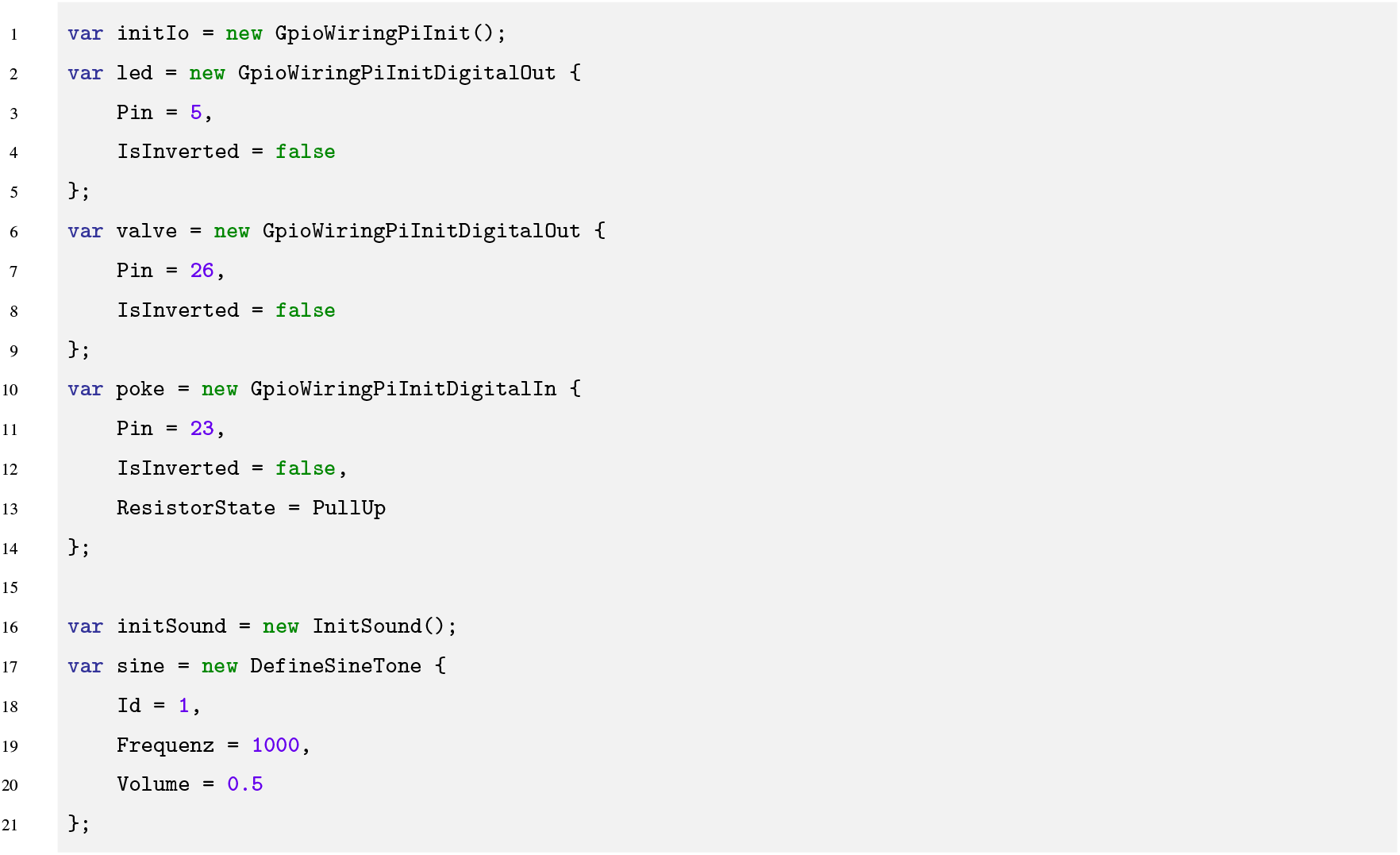

**Listing 1**. Each generated object represents an initialization message to be serialized with Protobuf and transmitted to LabNet. The first message initializes the digital I/O interface with WiringPi pin notation. The next three initialize an LED, a valve, and a poke sensor on the WiringPi interface. The fifth creates the sound generator on the headphone jack. The last, initializes a sine tone with 1kH frequency and 50% volume. Object initialisation in C# notation. Serialization and TCP/IP communication not listed.

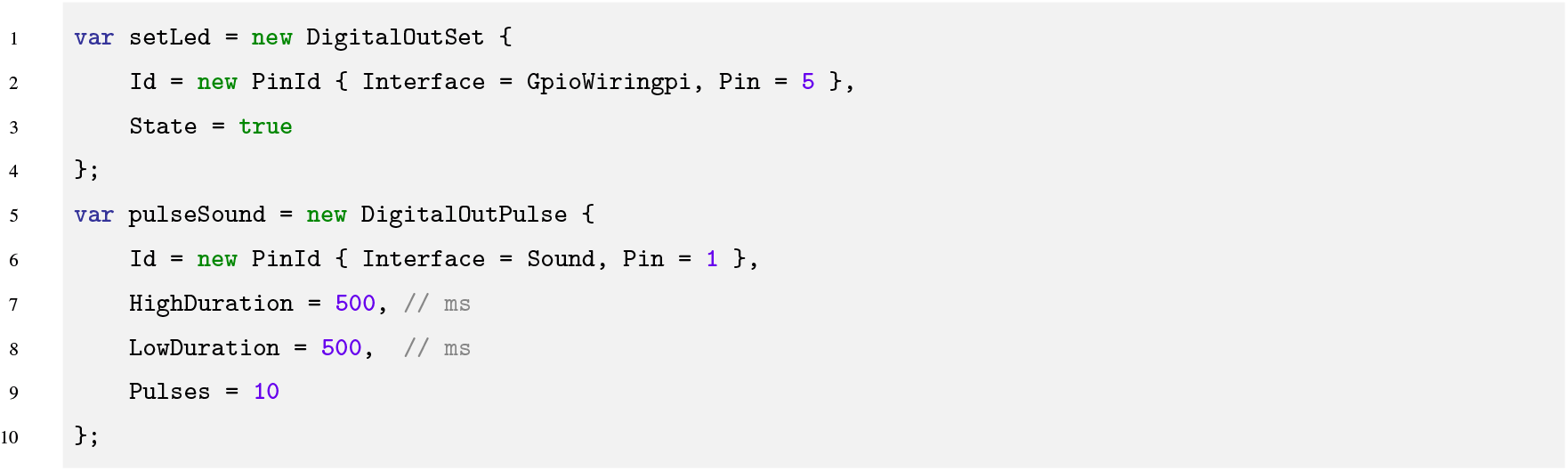

**Listing 2**. Information for transmission via Protobuf, building on the initialization from Listing 1. The LED is set to ON state (until an OFF command). A sound, defined as 1 kHz in Listing 1, is emitted as 10 pulses of 500 ms. Object initialisation code in C# notation.

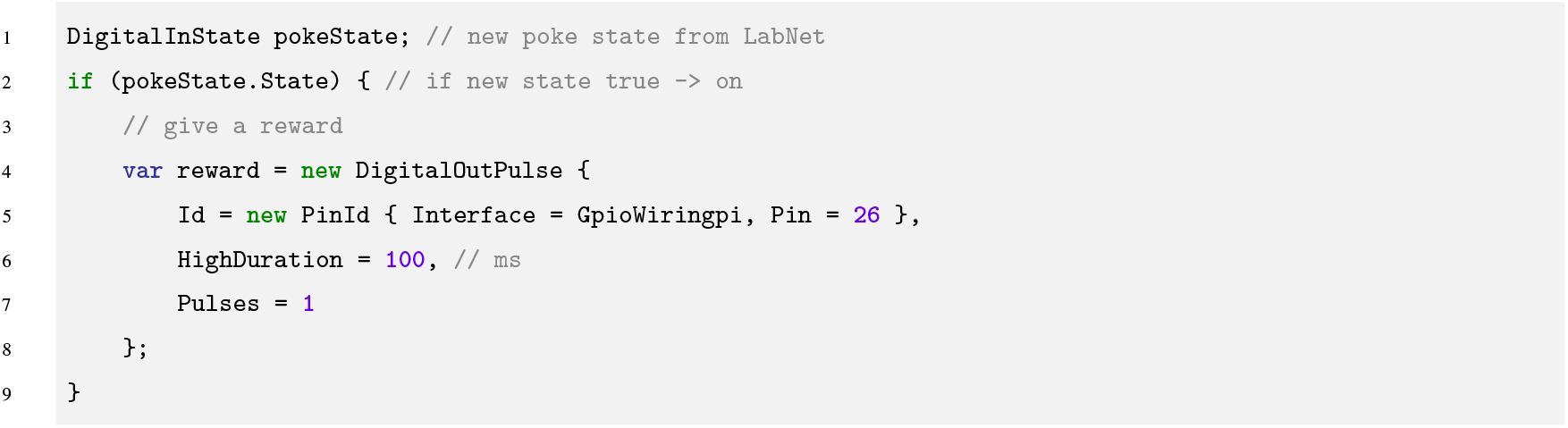

**Listing 3**. In this example, LabNet has transmitted via Protobfuf a new poke sensor state (pokeState) to the PC. If the new poke state is true, a new message directs LabNet to deliver a reward by opening valve at pin 26 for 100 ms, as initialized in Listing 1. Code shown in C# notation.

In Listing 3, a liquid reward valve is opened for 100 ms.

A typical experiment in combination with LabNet comprises several phases:

- establishing a TCP/IP connection;
- initializing all hardware components;
- turning stimuli on or off in a specific order;
- waiting for an animal reaction and potentially providing a reward.

## Related work

Most of the comparable software control tools published for behavioural experiments are more general packages. In addition to hardware control, they offer a more or less powerful tool for creating experiments, a user interface and a possibility to visualize the data. Although LabNet is only responsible for the hardware, a comparison is still worthwhile.

### Wisker-Server

The development of Whisker (***Cardinal and Aitken, 2010***) control suite started in 1999 by Cardinal and Aitken at the Department of Experimental Psychology, University of Cambridge and is ongoing. Initially, the aim was to use the existing resources of a PC and plugged-in IO cards to control behavioural experiments with visual stimuli and touch screens in several boxes simultaneously. This was solved by an additional software layer where Whisker operates as the server and controls the hardware. The clients must connect to the server over TCP/IP, and each one controls an experiment in one of the chambers. The clients themselves can be written in any programming language. Communication occurs through a plain-text protocol.

Because of the outsourcing of the experiments to the clients, Whisker’s approach is similar to ours. Due to the flexibility in implementing the clients, complex experiments can be realized with Whisker. Hardware support includes digital I/O devices (National Instruments, Advantech, etc.), visual stimuli on computer monitors, touchscreens, audio, and more. Whisker is commercially used in “ABET II” by Campden Instruments Ltd.

### pyControl

pyControl (***Akam et al., 2022***) is an open-source hardware and software framework for controlling behavioural experiments. The hardware is based on the MicroPython microcontroller that typically control a single experimental box each. Several pyControl breakout boards can connect to a PC via USB. Each board has six so-called behaviour ports and four BNC ports. Each port can be connected to a module: to drive LEDs, nose-poke sensors, stepper motors and speakers. Two behaviour ports have I2C internally and can drive a port expander module to increase the number of ports.

Tasks on the MicroPython microcontroller and pyControl on the PC use Python. A task is defined as a finite-state machine. It comprises a collection of states and events that cause the switch between states. In data management, all events and state changes are stored with timestamps.

pyControl provides sufficient I/O ports to realize most tasks on a system. However, for the hardware types, we are limited to the firmware capabilities and available modules, although free wiring is also possible. The mandatory requirement to define the task as a state machine can be useful but may also become a limitation.

### Bpod

Bpod (***Sanders, 2021***) was originally developed in the Brody lab and is now maintained by Josh Sanders (Sanworks LLC.). It has been also expanded to PyBpod as a python port of the Bpod Matlab project by members of the Champalimaud Foundation. Bpod offers only four I/O ports but has additional module ports that each provide an interface to Arduino-powered modules. Thus, Bpod gains additional flexibility: analog I/O, I2C, Ethernet, and more can be accessed via these modules.

A MATLAB package is offered to write experimental tasks. Unfortunately, the package documentation is limited. The tasks are also defined as finite-state machines. After starting the task, the state machine is transferred to the Bpod. From there, it communicates with the MATLAB frontend. This design results in the restriction that only a single Bpod can be controlled per MATLAB session. Therefore, Bpod is much more limited regarding software than pyControl or Whisker. Multiple systems cannot run simultaneously, and the functionality is limited by the firmware and the state machine.

### Autopilot

Autopilot ***Saunders and Wehr*** (***2019***) is an open-source framework for behavioural experiments developed in the Wehr Lab at the University of Oregon. It uses Python, and the target platform is the Raspberry Pi.

The focus of Autopilot from the beginning has been the ability to control multiple systems. The basic unit in the software architecture of Autopilot is an agent. Each agent runs on its own Raspberry Pi and can communicate with other agents. Currently, three types of agents exist: **terminal, pilot** and **child**.

**Terminal** agents are the only user-oriented with a graphical user interface. They are responsible for data logging and visualization. The experimental tasks are also managed here and transferred to the **pilots**, that are also responsible for experimental task execution. The pilots communicate with the external hardware that is connected to the Raspberry Pi and forward the experimental data to the **terminals** for logging or visualization. Each **pilot** can also have several **child** agents. **Child** agents can take over a part of a task if the task has been configured accordingly. The child agents are invisible to the terminals and communicate only with their parent **pilot**.

Among all tools discussed here, Autopilot offers most flexibility. It already supports a whole range of hardware. This includes digital I/O, audio, cameras and some sensors such as temperature. Moreover, since it is open-source, support for additional hardware can be added. New behavioural experiments can also be implemented. However, in both cases, we are limited to Python. The performance of Autopilot is good. However, all other tools including LabNet offer better performance as Autopilot (see Performance Evaluation).

## Performance Evaluation

Because the neurons in the brain work in the millisecond range, the response times in behavioural experiments are critical and should match that range.

LabNet was subjected to three tests to determine the latency time when executing different commands. A RasberryPi was connected to a PC via a router. Each test was run 100,000 times. For all tests, the client ran on a Windows PC (Intel Core i7-6700 3.4 GHz with 16 GB RAM). To allow a comparison, the client was implemented in three languages: Python, C#, and C++. We used Python version 3.8. Python tests ran directly on top of the socket, synchronously with no additional software layers. In Python, it is also essential to deactivate Nagle’s algorithm. The C# version was implemented under .NET 4.8 and used Akka.NET, an actor framework. For C++, we used Visual Studio 2019, Boost version 1.75 and SObjectizer 5.7.2. Thus, all tests in C# and C++ were implemented as actors inside an actor framework. The source code of all tests is included in the GitHub repository under “examples”.

LabNet was also compiled with different optimizations to investigate the performance effects of the different RasberryPi boards. However, GCC 8.3 was used for all versions. The first version had only the default release optimizations from CMake and could run on all RasPi boards. The optimization flags were as follows:

- `-mcpu=cortex-a7` for Pi 2;
- `-mcpu=cortex-a53` for Pi 3;
- `-mcpu=cortex-a72` for Pi 4;
- `-mfpu=neon-vfpv4 -mfloat-abi=hard` - floating-point number optimizations for all versions.

The influence of experimental data processing on the latency was not considered in the test runs. However, C# and C++ already used an actor framework for asynchronous message processing and both thus simulated the processes of an experiment in a first approximation. In Python all tests ran synchronously.

### ID test

In the “id test”, the client only requests the ID of a LabNet server, without any operations to the peripheral hardware. It measures the time interval between sending the request and arrival of the answer. As only message handling and network communication is measured it is a type of ping measurement.

The message from the client has only 2 bytes. The server response is 12 bytes long and contains a short `string` with two major and minor version numbers.

The results are shown in Figure 3a. All responses were sent within one millisecond. All three models are significantly different, where Raspi2 is the slowest and Raspi4 is the fastest. For C# the median on the RasPi 4 is 0.36 ms and 0.97 ms on the RasPi 2.

**Figure 3.**
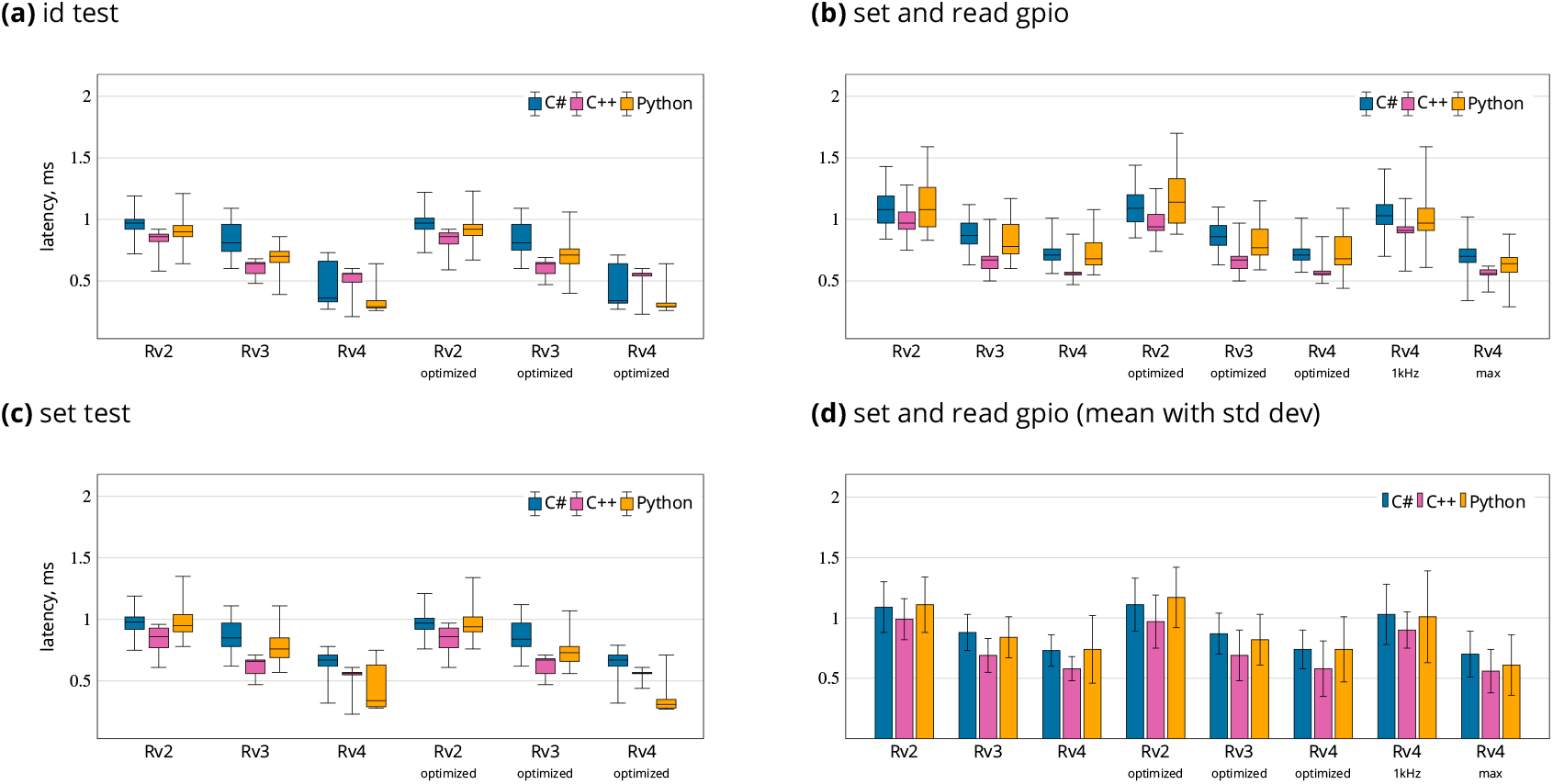
Results from LabNet performance tests: (a) id test as a Ping equivalent, (c) setting one output pin, (b, d) setting one output pin and detecting state change from a connected input pin. Tests were repeated 100,000 times and results are in milliseconds. Tests were performed on three different RaspberryPi boards: Rv2 is 2B, Rv3 is 3B+ and Rv4 is 4B with 1GB RAM. *optimized* refers to the LabNet with some additional optimization flags. 1kHz refers to the version without optimisations running on Rv4 with 1kHz polling and *max* to non-stop polling. Box plots in a to c show median, lower and upper quartile, and whiskers the 2.5th and 97.5th percentiles. Data in (d) given as means and STD.

### Set digital out test

In the “set out test”, a digital output is alternately set to 0 or 1. After the command for setting the pin has been received and processed, LabNet automatically sends back an acknowledgement. In this test, the time between sending the set command and receiving this confirmation was measured.

The set command from the client has 10 bytes. The server response is 22 bytes long, and includes the execution timestamp.

The results are shown in Figure 3c. The median was between 0.67 ms for 4 and 0.98 for 2, also for C#. Thus, switching a pin only increase latency of RasPi 4. But this is due to C#, as C++ and Python do not show this latency increase in comparison to the “id test”.

### Set and read GPIO

For this test, two IO pins of the RasPi were connected. One pin was defined as output and the other as input. The output was again alternately set to 0 or 1. We then measured the time interval between sending the output request and receiving the input status change. This approximates one of the most common situations in a real experiment with an animal. The only difference is that the animal has first to trigger the input and this leads to an output state change.

The client message has 10 bytes. The server sends this time two messages back. The first is the digital output state change confirmation. The second is the new digital input state. Both messages are 22 bytes long, including the execution timestamps.

Because this is the more important latency test, the results are summarized in Figures 3b and 3d. Pi 3 and 4 are similar. The median was 0.71 ms for 4 and 1.08 for 2. The 97.5 percentile was 1.01 ms for 4 and 1.43 ms for 2 and mean values are 0.73 ± 0.11 ms for 4 and 1.09 ± 0.21 for 2. All values for C#.

LabNet use a polling mechanism to detecting changes in digital inputs. By default, LabNet runs at 4kHz polling. But we also evaluated latencies for 1kHz and non-stop polling on Pi 4. Mean value in case of 1kHz is 1.03 ± 0.25 ms and 0.70 ± 0.19 ms for non-stop. Thus, 4kHz offers practically identical results compared to non-stop polling, but only utilizes 10% of the CPU.

The compiler optimization flags did not influence any of the tests. This indicates that LabNet has no performance issues on the RasPi.

The 1 GigE update from the RasPi 3B+ onwards causes a performance increase over model 2, although it is only with model 4 that the network controller is no longer integrated via USB.

### Comparison

Our comparison of LabNet latency performance with other software tools is summarized in Figure 4. Different from LabNet all of the other tools run locally and do not send commands over the wire in the network.

**Figure 4.**
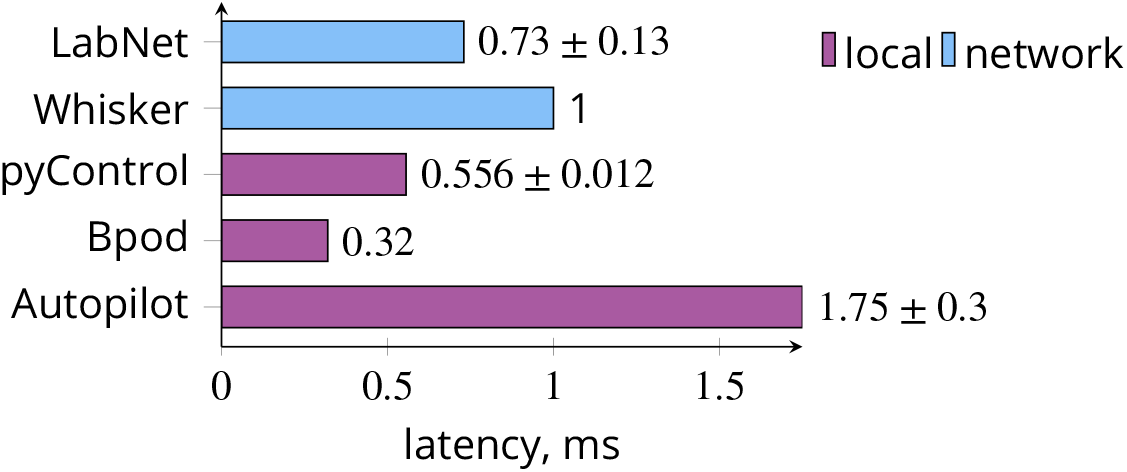
Comparison of execution latencies. Whisker server implements a 1 kHz polling frequency on the PC. LabNet for digital input polling depends on internal 4 kHz polling frequency. Only for LabNet do the latencies include the message transfer over the ethernet wire. Values give means with STD. LabNet value is for C# client and Raspi 4.

In Whisker, the communication occurs over the network; however, both Whisker and all task clients usually run on the same PC. Such communication is extremely fast and is also reported by the Whisker authors (***Cardinal and Aitken, 2010***) to require only 0.066 ms. The 1 ms latency comes from Whisker’s internal 1 kHz polling frequency for processing incoming commands.

Autopilot runs in a Rasberry Pi swarm. However, the experiments for which the latencies are reported ran locally. In the test, the time between detection of a photo gate interruption and the start of sound output via the headphone jack was 1.75 ± 0.3 ms. The latency times over the network cause additional delay (***Saunders and Wehr, 2019***). Autopilot requires 4.9 ± 0.47 ms from sending to receiving a message without a payload. In a real communication with a response, this value would double and add to execution time. That is, in the Set and read GPIO, Autopilot would require approximately 12 ms, and in the Set digital out test, it would require approximately 7 ms. The pyControl latency is 556 ± 17 µs (***Akam et al., 2022***) from detecting the change on a digital input to updating the state of a digital output. The pyBoard run under the same “low load” conditions as LabNet, i.e. only two pins were handled.

Bpod latency is 0.22 ms from detecting the trigger for a sound playback by the Bpod finite state machine until the sound output begins plus 0.1 ms to detect a change on a digital input. The value was published on the official Bpod site.

According to these measurements, LabNet achieves latency times comparable to or better than locally executed applications, even despite controlling the hardware over the network via TCP/IP.

## Conclusion and Outlook

LabNet provides a software platform to control the hardware in laboratory experiments, and was used by us in behavioural experiments. Through the combination of C++, SObjectizer and Protobuf, LabNet achieves almost real-time hardware control via network communication. By using the actor framework SObjectizer, it is relatively easy to add new functionality despite programming close to the hardware with C++. The choice of the Raspberry Pi as the hardware platform allows experimental setups to connect a wide variety of readily available low-cost sensors and actuators.

Already with this initial version, a whole range of hardware modules and interfaces can be addressed with LabNet via network. For example, GPIOs, communication via the UARTs, sound output via a headphone jack or HDMI and some Raspberry Pi HATs developed in-house are supported. This alone allows many types of experiments. With hardware adaptors for the Raspberry Pi the experimental control can extend to the wide range of available modules for operant experiments from open source Bpod and pyControl, but also to the commercial systems of MedAssociates, Coulbourn Instruments, and Lafayette Instruments. Visual stimuli, touchscreen support and communication via I2C will be included in the next version. Support for a Raspberry Pi based configuration file is also planned. This would make configuration via the network no longer necessary and LabNet could already initialize the hardware correctly on Raspberry Pi start.

One goal on our roadmap is to implement a software plug-in system. This will make it possible to support new hardware without LabNet recompiling. This will require a rework of the current LabNet API. In addition, the message protocol will be modified so that messages unknown at compile time can be received and sent.

## Code availability

The source code of LabNet is available over the GitHub repository under the GPL-3.0 license and the version used here is available over the Zenodo archive: https://doi.org/10.5281/zenodo.5549518.

Data of the performance measurements together with the source code for the graphs are also accessible via the GitHub repository.

There are instructions for two possible compilation paths. The first is on the Raspberry Pi with Visual Studio Code and CMake. The second is with Visual Studio 2019 and Docker. The archive also contains the source code of all tests from Performance Evaluation under “examples”.

## Acknowledgements

Support for this work was received through the Deutsche Forschungsgemeinschaft (DFG, German Research Foundation), SFB 1315, project-ID 327654276, and EXC 257: NeuroCure, project-ID 39052203.

## Performance tests

**Appendix 0 Table 1.**
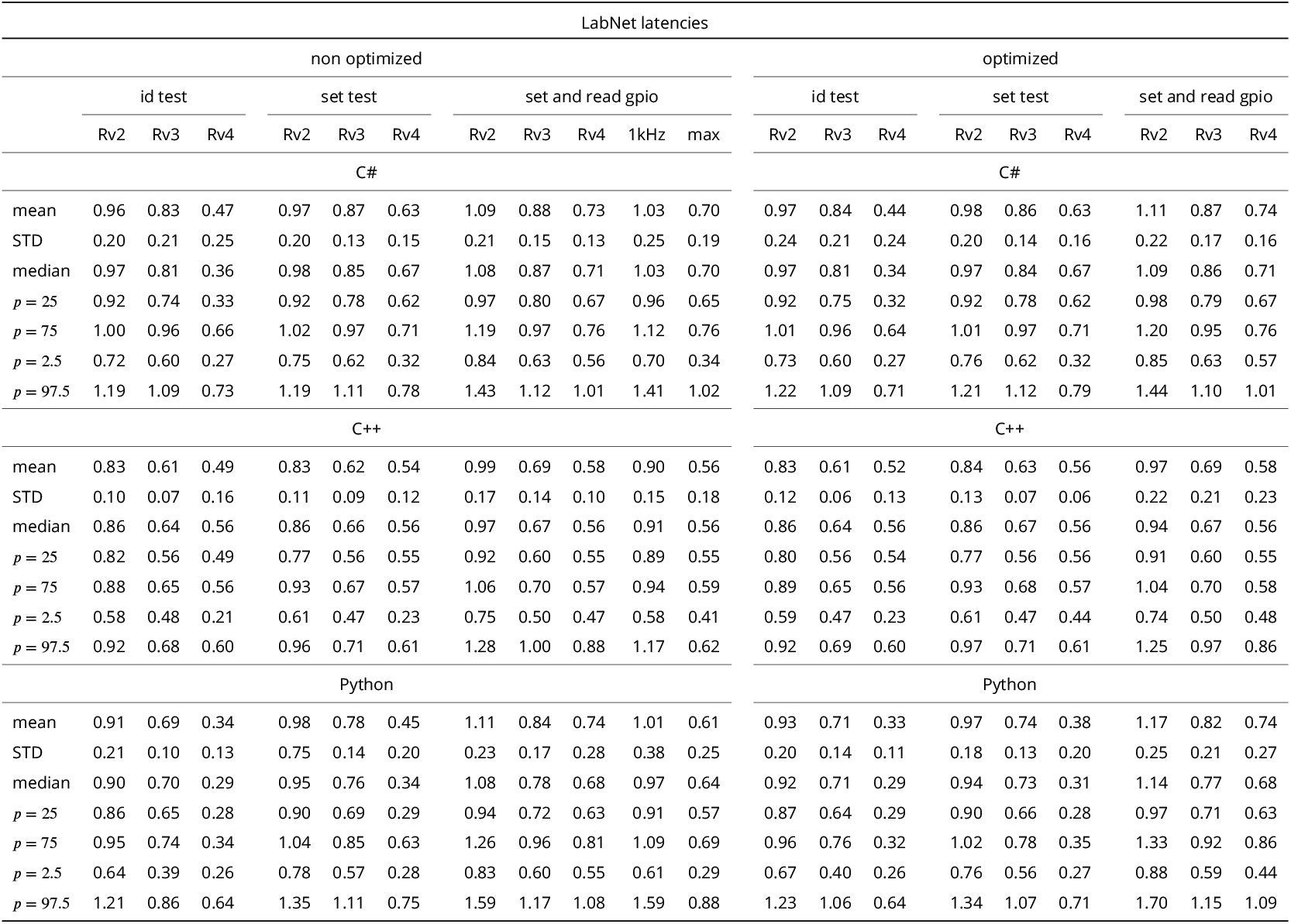
The results of all three LabNet performance tests from section Performance Evaluation. Each test was called 10000 times. The results are in milliseconds. *p* is percentile, median is also *p* = 50. All tests were performed on three different RaspberryPi boards: Rv2 is 2B, Rv3 is 3B+ and Rv4 is 4B with 1GB RAM.

## Footnotes

1 All images and diagrams are generated with Ti*k*Z. The mouse image is from Ti*k*Zlings package and the PC image from mo pTi*k*Z.

